# Limited recognition of *Mycobacterium tuberculosis*-infected macrophages by polyclonal CD4 and CD8 T cells from the lungs of infected mice

**DOI:** 10.1101/697805

**Authors:** Yash R. Patankar, Rujapak Sutiwisesak, Shayla Boyce, Rocky Lai, Cecilia S. Lindestam Arlehamn, Alessandro Sette, Samuel M. Behar

**Author notes:** Correspondence: Samuel M. Behar, E-mail address (SMB).

## Abstract

Immune responses following *Mycobacterium tuberculosis* (Mtb) infection or vaccination are frequently assessed by measuring T cell recognition of crude Mtb antigens, recombinant proteins, or peptide epitopes. We previously showed that not all Mtb-specific T cells recognize Mtb-infected macrophages. Thus, an important question is what proportion of T cells elicited by Mtb infection recognize Mtb-infected macrophages. We answer this question by developing a modified elispot assay using viable Mtb-infected macrophages, a low multiplicity of infection and purified T cells. In C57BL/6 mice, CD4 and CD8 T cells were classically MHC restricted. Comparable frequencies of T cells that recognize Mtb-infected macrophages were determined using interferon-γ elispot and intracellular cytokine staining, and lung CD4 T cells more sensitively recognized Mtb-infected macrophages than lung CD8 T cells. Compared to the numbers of Mtb antigen-specific T cells for antigens such as ESAT-6 and TB10.4, low frequencies of pulmonary CD4 and CD8 T cells elicited by aerosolized Mtb infection recognize Mtb-infected macrophages. Finally, we demonstrate that BCG vaccination elicits T cells that recognize Mtb-infected macrophages. We propose that the frequency of T cells that recognize infected macrophages could correlate with protective immunity and may be an alternative approach to measuring T cell responses to Mtb antigens.

## Introduction

The WHO estimates that 23% of the world’s population is latently infected with *Mycobacterium tuberculosis* (Mtb), the causative agent of tuberculosis (TB), and 10 million active cases are reported every year^1^. An incomplete understanding of the host-pathogen interactions and the lack of known correlates of protective immunity have hampered the development of an efficacious TB vaccine.

Mtb infection elicits CD4 and CD8 T cell responses in both humans and animal models, and their role in immunity to primary disease is widely appreciated. Numerous vaccine strategies use immunodominant antigens to elicit T cell responses. Most human and murine vaccine studies rely on using crude Mtb fractions or Mtb peptides as antigens to assess the immunogenicity and function of vaccine-elicited T cells. An underlying assumption has been that most Mtb antigen-specific T cells elicited during natural infection will recognize Mtb-infected antigen presenting cells (APC). However, the parameters used to measure vaccine immunogenicity such as cell numbers or cytokine responses of antigen-specific T cells after stimulation with antigen have not correlated with, or predicted the protective potential of vaccines^2, 3^.

Recent data challenge the assumption that all Mtb-antigen specific T cells primed following infection recognize Mtb-infected macrophages. We find that CD8 T cells specific for TB10.4_4-11_, an immunodominant epitope in C57BL/6 mice, do not recognize Mtb-infected macrophages and vaccination with TB10.4_4-11_ does not confer protection^4, 5^. Other studies find that CD4 T cells specific for Ag85b_240-254_, another immunodominant antigen, have a weak response in granulomas due to limited local antigen presentation by infected myeloid cells ^6, 7^. Yet optimal control of Mtb *in vivo* requires direct recognition of infected myeloid cells by CD4 T cells^8^. The chief paradigm of T cell-based vaccines is that the elicited T cells must recognize Mtb-infected macrophages to confer protection.

It is difficult to reconcile the profound immunodominance of some Mtb antigens with the failure of T cells specific for those antigens to recognize Mtb-infected macrophages^4^. Importantly, following aerosol infection, Mtb disseminates to the mediastinal lymph node, where T cells are first primed by dendritic cells, which then expand and traffic to the lung^9, 10^. We speculate that there may be a mismatch in the antigens presented (or cross-presented) by uninfected DC in the lymph nodes and antigens presented by infected macrophages in the lung. Thus, T cells primed in the lymph nodes during natural infection may not necessarily recognize antigens presented by Mtb-infected macrophages in the lung^11^. Regardless of the mechanism, we wondered whether the inability of some T cells to recognize Mtb-infected macrophages might explain why the number of antigen-specific T cells may not necessarily correlate with vaccine-induced protection.

To assess T cell recognition of Mtb-infected macrophages we developed a modified elispot assay based on interferon (IFN)-γ spot forming cells (SFC). Using a low multiplicity of infection (MOI), we quantify the frequency of T cells that recognize Mtb-infected macrophages during primary infection in mice. We find that an unexpectedly low frequency of *ex vivo* CD8 and CD4 T cells recognizes Mtb-infected macrophages. We demonstrate that majority of the T cells from C57BL/6 mice that recognize Mtb-infected macrophages are conventionally MHC-restricted T cells. Our data shows that CD4 T cells efficiently detect Mtb-infected macrophages at a lower MOI, whereas CD8 T cells only recognize more heavily infected cells. Using proof-of-concept vaccination studies, we show that BCG elicits T cells that recognize Mtb-infected macrophages. We envision this novel assay as a complementary approach to immunogenicity studies and mycobacterial growth inhibition assays. By specifically measuring the frequency of vaccine-elicited T cells that recognize Mtb-infected macrophages pre-challenge, this assay could provide another criterion to help screen and prioritize the selection of T cell-based vaccines for preclinical and clinical development.

## Results

### Measuring T cell recognition by the Mtb-infected macrophage elispot (MIME)

We modified our established *in vitro* macrophage infection model ^4^. We aimed to maximize the percentage of infected macrophages, preserve their viability, and achieve a physiologically relevant MOI ^12^. Using YFP-expressing H37Rv at a multiplicity of infection (MOI) of 4 to infect thioglycolate-elicited peritoneal macrophages (TG-PM), we found that 70% of macrophages were infected with 93% viability after 18-24 hours (Fig.1a and 1b). In contrast, fewer than 35% of macrophages were infected at an MOI of 1 (Fig.1c). Although 90% of the macrophages were infected at an MOI of 20, there was a drastic decrease in macrophage viability (Fig.1c). Using an MOI of 4 for our assays led to a median effective MOI of about 5 (range 2 – 13) (Fig.1d). Thus, an MOI of 4 maximized the percentage of infected macrophages while maintaining cell viability.

**Figure 1.**
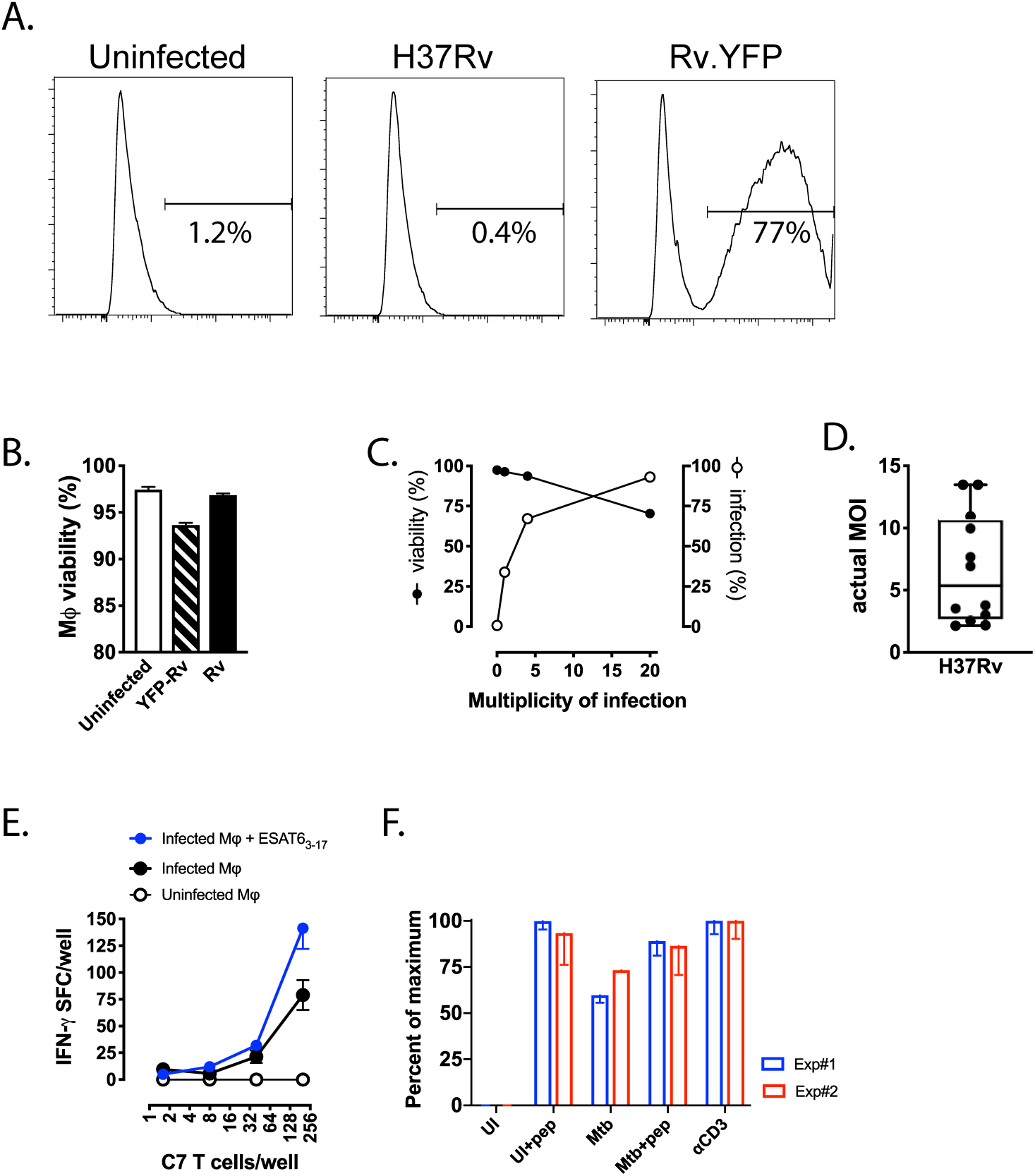
An assay to measure T cell recognition of Mtb-infected macrophages. A, B) After infection with H37Rv or Rv.YFP at an MOI of 4 for 18 h, CD11b^+^-enriched thioglycolate-elicited peritoneal macrophages (TG-PM) were assessed for the percentage of cells that were infected (A) and viable. C) TG-PM were infected with Rv.YFP at an MOI of 1, 4 or 20, for 18-24 h and the percentage of infected macrophages and cell viability were assessed. D) Macrophages were infected with H37Rv with an MOI of 4 for 18-24 h and the actual MOI was determined by plating CFU. E) The Mtb-infected macrophage ELISPOT (MIME) assay was performed by infecting macrophages with H37Rv at an MOI of 4, for 18-24 h. A titrated number of C7 T cells were mixed with polyclonal T cells (10^5^/well) from uninfected mice and added to Mtb-infected macrophages (10^5^/well). Where indicated, the ESAT-6_3-17_ peptide was added to the wells. The assay was performed as described in the “Methods”. F) The P25 T cell line was added to Mtb-infected macrophages, and the MIME assay performed. UI, uninfected macrophages; Mtb, Mtb-infected macrophages; pep, Ag85b_240-_ 254 peptide; αCD3, soluble anti-CD3 mAb. Two independent experiments are shown. Each experiment was normalized by subtracting the background and defining the αCD3 response as the maximal (i.e., 100%) response. Data are representative of 2 independent experiments with 3 technical replicates per experiment that yielded similar results (A, B, C, E, F), or 12 independent experiments with 3 technical replicates (D).

We next established an IFN-γ enzyme-linked immunospot (elispot) assay to measure the frequency of T cells that recognize Mtb-infected macrophages by culturing purified T cells with Mtb-infected macrophages. To first validate the assay, we diluted ESAT-6-specific CD4 T cells (hereafter, C7 T cells; see methods) into an excess of splenic T cells from uninfected C57BL/6 mice. Minimal activation of C7 T cells occurred when they were cultured with uninfected macrophages (Fig.1e). There was no activation of the naïve C57BL/6 T cells when cultured with infected macrophages or uninfected macrophages and ESAT-6_3-17_ peptide (data not shown). When the C7 and naïve T cell mixture was cocultured with Mtb-infected macrophages, specific spots were produced by 40-50% of the input C7 cells, and specific spots were detected even when their frequency was as low as 0.04% of the total T cells (Fig.1e). When ESAT-6_3-17_ peptide was provided in excess, 70-80% of C7 T cells produced IFN-γ (Fig.1e). We then used Ag85b-specific CD4 T cells (hereafter, P25 cells) in this assay. We observed that more than half of the P25 T cells that recognized Ag85b_240-254_ peptide could recognize Mtb-infected macrophages (Fig.1f). Furthermore, infected macrophages did not inhibit T cell production of IFN-γ in response to the cognate peptide. Based on our results with C7 and P25 T cells, the <u>M</u>tb-Infected <u>M</u>acrophage <u>E</u>LISPOT (MIME) appeared to represent a sensitive and specific way to identify T cells that recognize Mtb-infected macrophages.

### A low frequency of T cells recognizes Mtb-infected macrophages

We next determined the frequency of polyclonal T cells from low-dose aerosol Mtb-infected mice that recognized Mtb-infected macrophages using the MIME. Among highly purified splenic CD4 and CD8 T cells from mice that had been infected for 4 or 8 wpi, a small population of splenic CD4 and CD8 T cells recognized Mtb-infected macrophages (Fig.2a). A negligible number of these T cells recognized uninfected macrophages; however, there was some activation of T cells from uninfected mice after culture with Mtb-infected macrophages. Based on these data, we calculated the frequency of splenic CD4 and CD8 T cells that recognized Mtb-infected macrophages as 2-3%, and 0.5-1%, respectively, between 4-10 wpi (Fig.2b).

**Figure 2.**
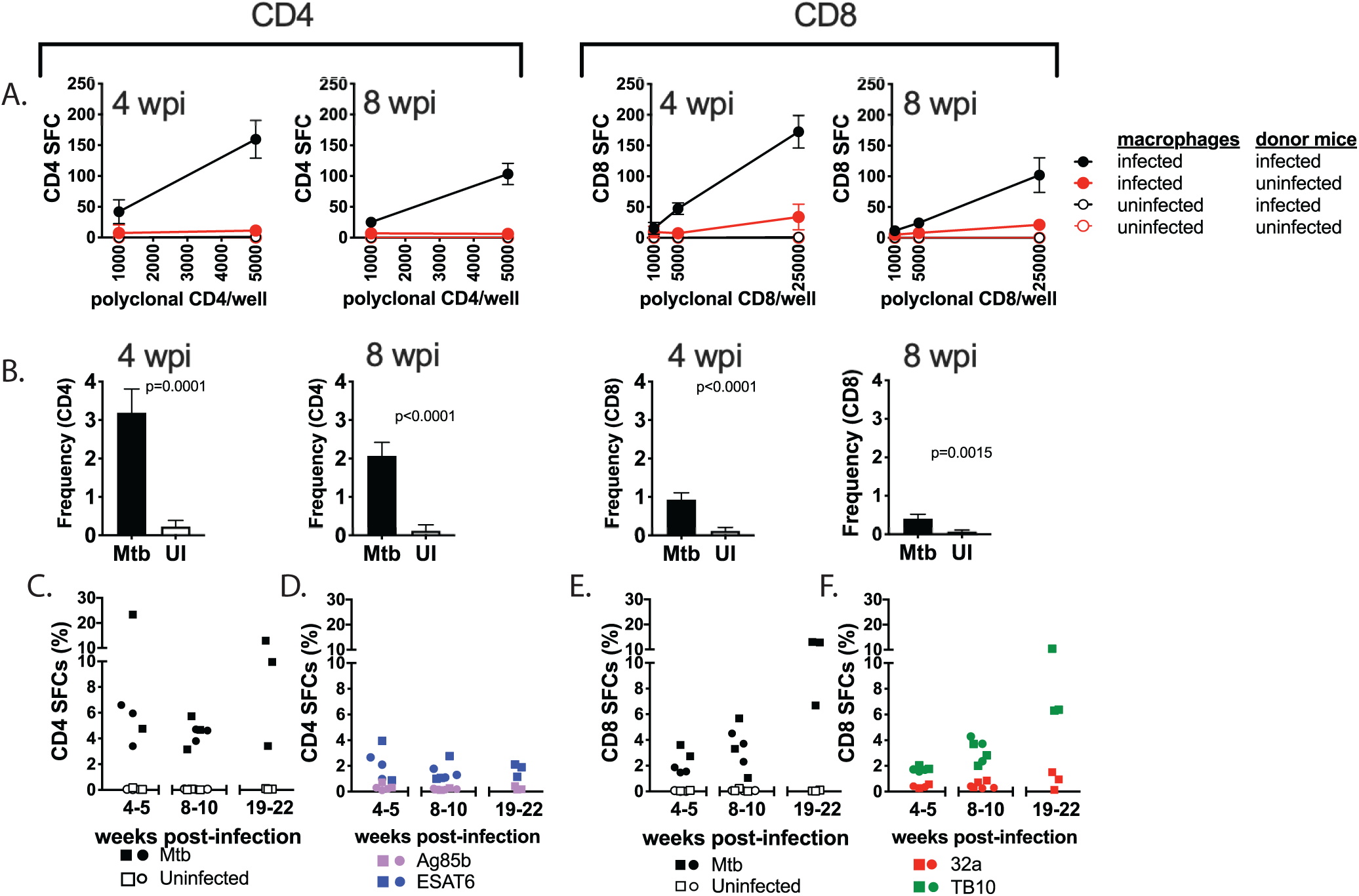
A low frequency of T cells from infected mice recognize Mtb-infected macrophages. A) Splenic CD4 or CD8 T cells from mice infected for 4 or 8 weeks were added to Mtb-infected macrophages and the MIME assay performed. Each timepoint is the average of 2 independent experiments using pooled T cell samples from 2-3 mice per time point analyzed in triplicates. Black symbols, T cells from infected mice; red symbols, T cells from uninfected mice. Closed symbols, T cells cultured with Mtb-infected macrophages; open symbols, T cells cultured with uninfected macrophages. SFC, spot forming cell. B) Frequencies of T cells recognizing Mtb infected macrophages, as calculated from the MIME assay after subtracting the background (i.e., Mtb-infected macrophages alone). Mtb, T cells from infected mice; UI, T cells from uninfected mice. C, D, E, F) The MIME assay was performed using lung CD4 or CD8 T cells from mice infected for 4-5, 8-10, or 19-22 weeks, which were added to Mtb-infected macrophages or added to uninfected macrophages with the peptides indicated in the figure. The data are combined from 3-4 independent experiments, each with 2-3 technical replicates per timepoint. The squares denote pooled T cells samples from 2-4 mice, and the circles denote represent results from individual mice. Ordinary one-way ANOVA with Tukey’s multiple comparisons was performed by combining the results from individual and pooled mice. *, p ≤ 0.05.

A higher frequency of lung T cells recognized Mtb-infected macrophages (i.e., MIME^+^ T cells) compared to splenic T cells. Four to six percent lung CD4 T cells were MIME^+^ through 10 wpi and then trended to increase to 4-20% by 20 wpi (Fig.2c). The frequency of MIME^+^ CD4 T cells was higher than the combined frequency of CD4 T cells that made IFN-γ in response to Ag85b_240-254_ and ESAT-6_3-17_ (Fig.2d). Based on the known ability of ESAT-6-specific and Ag85b-specific T cell lines to recognize Mtb-infected macrophages (Fig.1 and ^4, 13^), these T cell specificities likely constitute a subset of the CD4 T cells that recognize Mtb-infected macrophages during infection.

Similarly, between 4-10 wpi, the frequency of MIME^+^ CD8 T cells remained relatively constant at 2-6% and then increased by 20 wpi (Fig.2e). The frequency of CD8 T cells recognizing Mtb-infected macrophages was generally higher than the frequency of T cells that made IFN-γ in response to 32a_93-102_ and TB10.4_4-11_ epitopes (Fig.2f). Although the frequency of TB10.4_4-11_-specific CD8 T cells tracked the frequency of MIME^+^ CD8 T cells, our prior work shows that TB10_4-11_-specific CD8 T cells do not recognize Mtb-infected macrophages^4^. In conclusion, although the frequencies of CD4 and CD8 T cells that recognize Mtb-infected macrophages increased over time, it was still a low percentage of the total splenic or pulmonary T cells.

### Lung T cells recognize Mtb-infected macrophages by a MHC-dependent manner

T cell activation results from TCR-mediated recognition of MHC-presented peptides (i.e., cognate activation) or from cytokine-driven stimulation by a TCR-independent mechanism (i.e., noncognate activation)^14^. IL-12, which is produced by Mtb-infected macrophages, is a principal driver of noncognate activation^15^. To discriminate between cognate and noncognate T cell activation, pulmonary CD8 T cells were cultured with Mtb-infected WT (i.e., MHCI-sufficient) or K^b^D^b−/−^ (i.e., MHCI^−/−^) macrophages. CD8 T cell recognition of Mtb-infected macrophages was largely MHCI-restricted since there was a 90% reduction in SFC in the absence of K^b^ and D^b^ (Fig.3a). Furthermore, blocking IL-12 had no effect on CD8 T cell recognition of Mtb-infected macrophages (Fig.3b).

**Figure 3.**
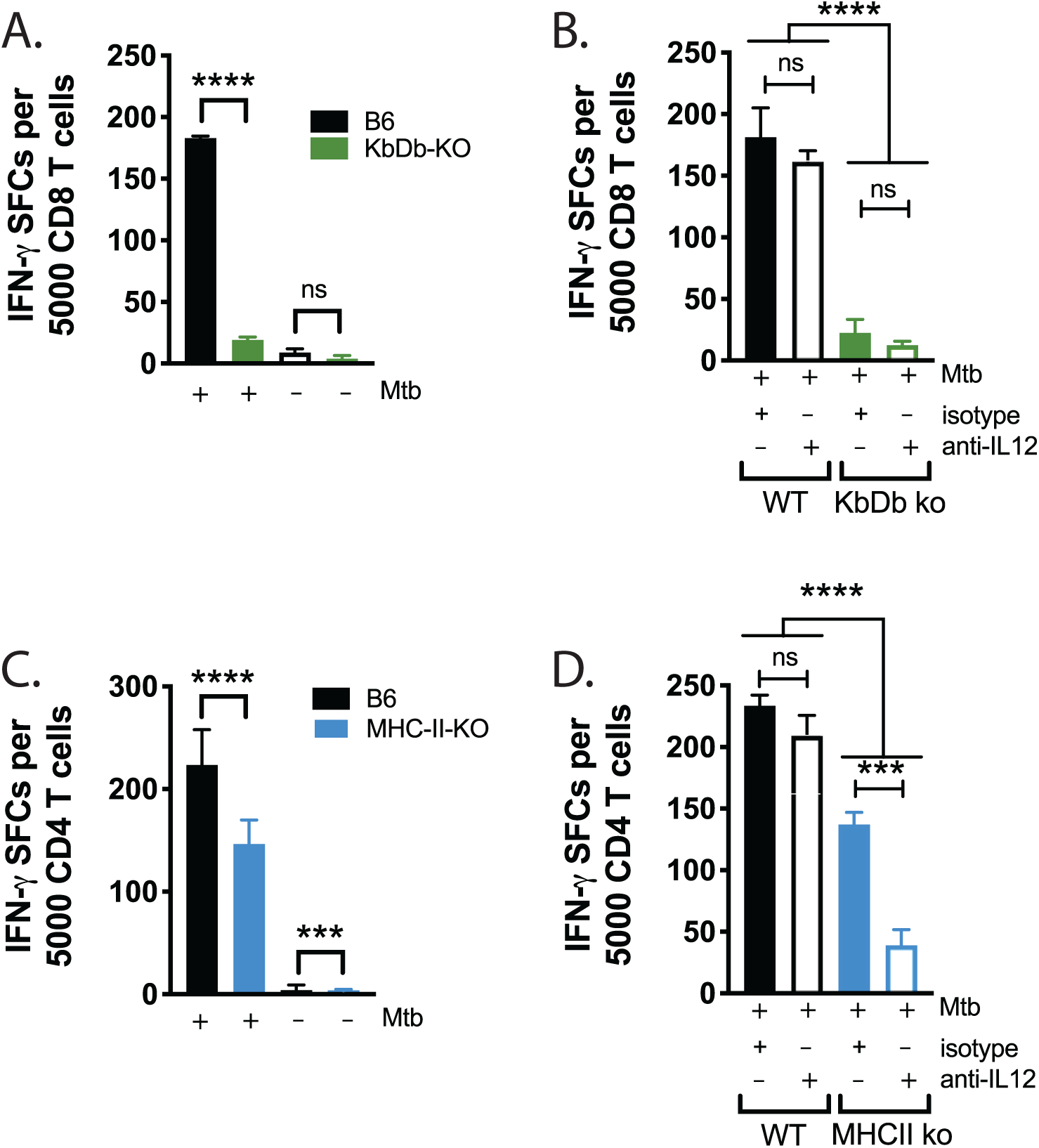
T cells that recognize Mtb-infected macrophages are MHC-restricted. (A, B) Pulmonary CD8 T cells from mice infected for 4-5 weeks were cultured with Mtb-infected WT or K^b^D^b−/−^ macrophages and the MIME assay performed. Where indicated, anti-IL12 blocking mAb or the appropriate isotype control antibody were added to the wells. (C, D) Pulmonary CD4 T cells from mice infected for 8-9 weeks were cultured with Mtb-infected WT or MHCII^−/−^ macrophages and the MIME assay performed. Blocking conditions as above. Data are representative of 2 independent experiments using pooled T cell samples from 2-4 mice per time point analyzed in 2-3 technical replicates. A one-way ANOVA with Tukey’s multiple comparisons was performed for statistical analyses, adjusted *p-*values: * ≤ 0.05, ** ≤ 0.01, *** ≤ 0.001, **** ≤ 0.0001.

To address the possibility of noncognate activation among CD4 T cells, we cultured pulmonary CD4 T cells with Mtb-infected WT (MHCII-sufficient) or *Ab1*^−/−^ (MHCII^−/−^) macrophages. Surprisingly, only a 40-50% reduction in SFC frequency was seen when CD4 T cells were cultured with Mtb-infected MHCII^−/−^ macrophages (Fig.3c). One possibility is that these CD4 T cells were nonconventional T cells^16, 17^; another possibility is that these T cells were conventional T cells, but in the absence of MHC, their IFN-γ production was driven by IL-12 produced by the infected macrophages^18, 19^. When CD4 T cells were cocultured with Mtb-infected WT macrophages in the presence of blocking antibodies to IL-12, no reduction in T cell activation was observed (Fig.3d). However, when CD4 T cells and MHCII^−/−^ Mtb-infected macrophages were cultured in the presence of blocking antibodies to IL-12, there was a significant reduction in the SFC compared to the control groups (Fig.3b). These data indicate that while IL-12 is sufficient for noncognate activation of lung CD4 T cells to secrete IFN-γ, the majority of the CD4 T cell-response is MHCII-restricted, and cognate activation of CD4 T cells does not require IL-12. In conclusion, our data show that majority of both CD8 and CD4 T cells recognize Mtb-infected macrophages through classical MHC recognition.

### T cell recognition of Mtb-infected macrophages measured by flow cytometry

We next adapted the MIME assay to a flow cytometry-based assay. Lung T cells from infected mice were used for the MIME assay and in parallel, were cultured with Mtb-infected macrophages and analyzed by intracellular cytokine staining (MIM-ICS). These two different assays led to similar estimates of the frequencies of CD4 and CD8 T cells that recognize Mtb-infected macrophages using IFN-γ as a readout (Fig.4a, 4b). Interestingly, while nearly all IFN-γ-producing CD8 T cells expressed CD69, a significant fraction of IFN-γ-producing CD4 T cells were CD69-negative (Fig.4a).

**Figure 4.**
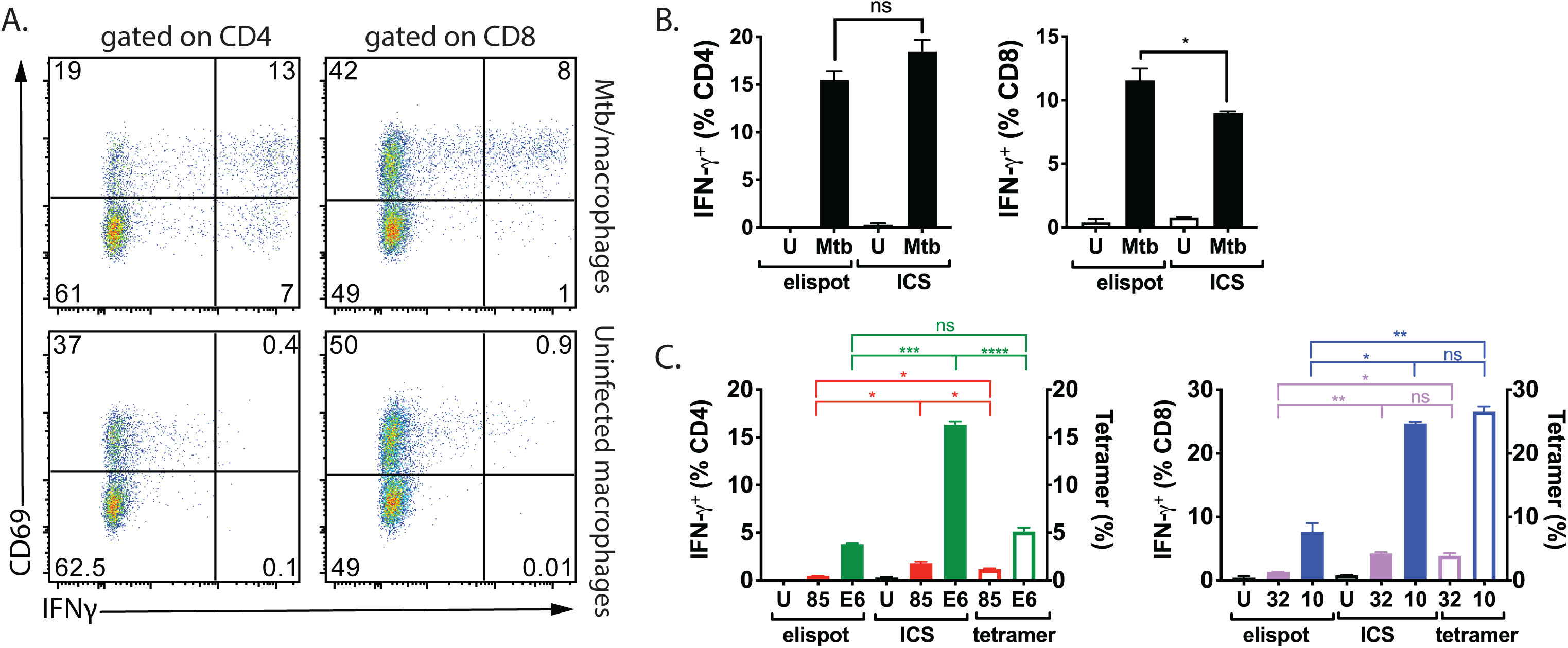
Similar frequencies of T cells recognizing Mtb-infected macrophages are detected by the MIME assay and MIM-ICS. (A) Pulmonary T cells were cultured with Mtb-infected macrophages and analyzed by ICS and flow cytometry. Representative flow plots showing the frequency of pulmonary CD4 (left column) or CD8 (right column) T cells expressing CD69 and producing IFN-γ after a 6 h culture with Mtb-infected macrophages. (B) The MIME or the Mtb-infected macrophage-ICS (MIM-ICS) assay were used to calculate the frequency of CD4 (left) or CD8 (right) pulmonary T cells that recognized Mtb-infected macrophages. U, uninfected macrophages; Mtb, Mtb infected macrophages. n.s., not significant (t-test). (C) The frequency of ESAT-6- or Ag85b-specific CD4 T cells (left) or the frequency of TB10.4- or 32a-specific CD8 T cells (right) among T cells from the lungs of Mtb-infected mice as determined by elispot or ICS, using peptide-pulsed uninfected macrophages, or tetramer staining. Data are representative of 2 independent experiments using pooled T cells from 5 mice at 5 WPI (shown, A – C) or 7.5 months post infection, analyzed in triplicates. Ordinary one-way ANOVA with Tukey’s multiple comparisons was performed for statistical analyses, adjusted *p-*values: * ≤ 0.05, ** ≤ 0.01, *** ≤ 0.001, **** ≤ 0.0001.

The frequencies of CD4 and CD8 T cells that recognized well-characterized class II and class I MHC-restricted Mtb epitopes were compared using elispot and ICS using uninfected macrophages, and tetramers. We detected 5.1% ESAT-6_3-17_/I-A^b^ tetramer^+^ and 1.2% Ag85b_240-254_/I-A^b^ tetramer^+^ CD4 T cells, consistent with published frequencies^7, 20^. After stimulation with the ESAT-6_3-17_ and Ag85b_240-254_ peptides the total frequency of CD4 T cells producing IFN-γ measured by ICS was greater than the frequencies determined using the tetramers (Fig.4c, left). Just as we observed a large population of CD69^−^ CD4 T cells that produced IFN-γ in response to Mtb-infected macrophages, CD69 was also absent from a large fraction of the CD4 T cells that produced IFN-γ after ESAT-6 stimulation (Fig.4c, left; Supplementary Figure 1). In contrast, the frequency of ESAT-6_3-_17- or Ag85b_240-254_-specific T cells measured by the elispot assay was lower than that determined by tetramer staining or by ICS (Fig.4c). For CD8 T cells, we detected 3.9% 32a_93-102_/K^b^ tetramer^+^ and 26.6% TB10.4_4-11_/K^b^ tetramer^+^, respectively, consistent with published frequencies^5, 21, 22^. In contrast to the results obtained for CD4 T cells, the frequency of CD8 T cells that recognized 32a_93-102_ and TB10.4_4-11_ based on ICS was similar to the frequency determined using tetramers (Fig.4c, right). Similar to the results obtained with CD4 T cells, the frequency of antigen-specific T cells determined by the elispot assay for 32a_93-102_ and TB10.4_4-11_, was lower than the frequency determined using ICS or tetramers.

### CD4 and CD8 T cells differ in their ability to recognize Mtb-infected macrophages

Hypothetically, the aggregate of T cell recognition of individual Mtb antigens should approach or be equivalent to the degree to which T cells recognize Mtb-infected macrophages. To assess whether T cell recognition of Mtb antigens is similar to the recognition of Mtb-infected macrophages, we took advantage of the megapool of 300 peptides (p300) representing 90 antigens, all of which are frequently recognized by human CD4 T cells from healthy IGRA+ individuals^23^. We cocultured lung cells obtained from mice that had been infected for four weeks with Mtb-infected macrophages, or with the p300 megapool, and measured IFN-γ production by T cells. The CD4 T cell response was skewed more towards the recognition of Mtb-infected macrophages than to the p300 megapool, indicating that some epitopes presented by Mtb-infected murine macrophages are not represented in the p300 megapool. In contrast, the CD8 T cell response was skewed towards the recognition of the p300 megapool (Fig.5a). Thus, CD8 T cells recognize Mtb antigens that are either poorly presented or are not presented by Mtb-infected cells but are represented in the p300 pool (e.g., TB10.4_1-15_). Compared to CD8 T cells, a greater frequency of CD4 T cells recognized Mtb-infected macrophages, indicating that CD4 T cells recognize Mtb-infected cells better than CD8 T cells at this time point (Fig.5a).

**Figure 5.**
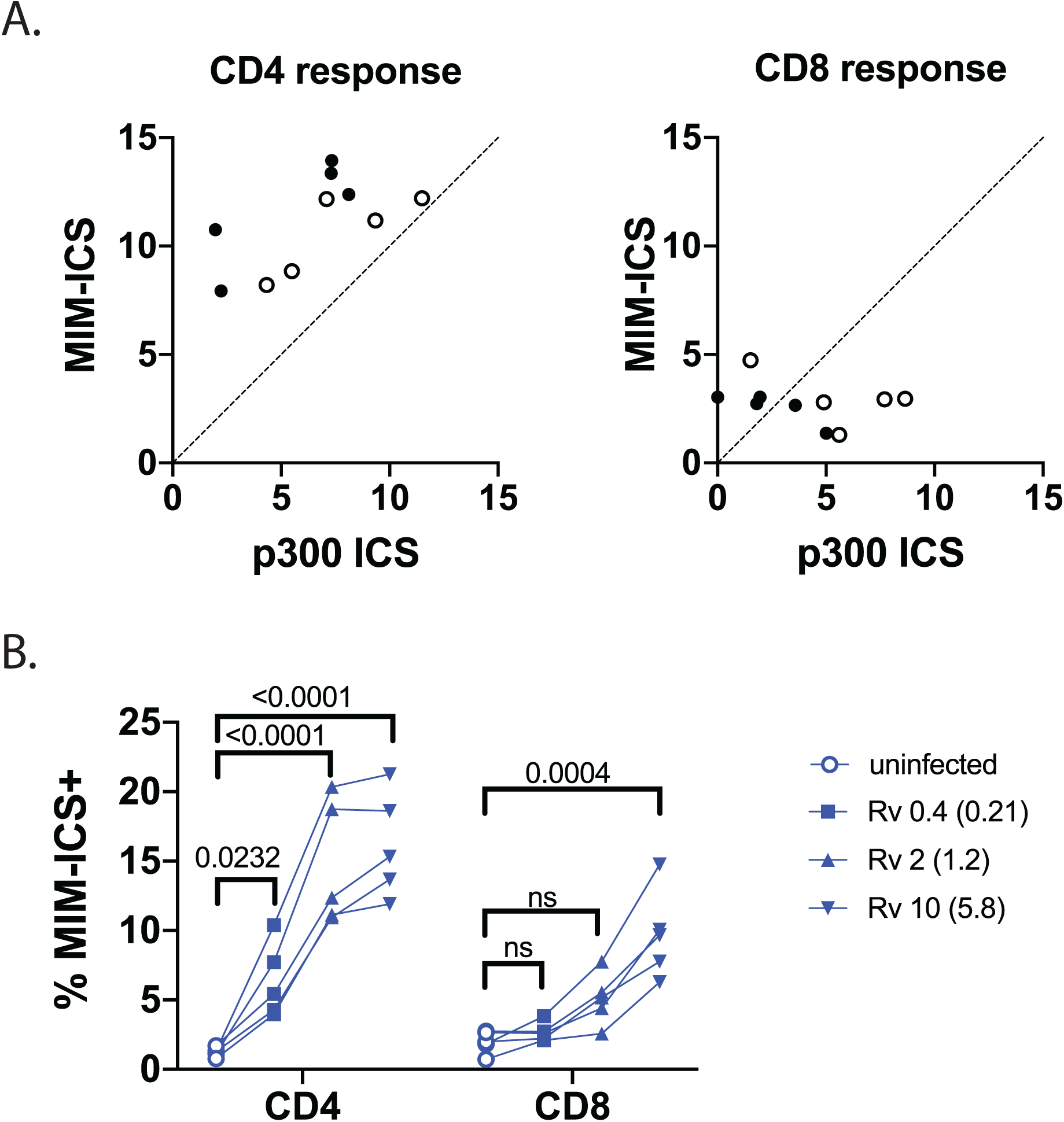
Differences in CD4 and CD8 T cell recognition of Mtb-infected macrophages. A) Lung cells obtained from Mtb-infected C57BL/6 mice and the CD4 (left) and CD8 (right) T cell recognition of Mtb-infected macrophages or the 300 peptide megapool (p300) was compared by ICS. Combined data from 2 independent experiments (identified by open or closed symbols), each with 5 mice/group, analyzed 4 wpi. B) The frequency of pulmonary CD4 and CD8 T cells that produced IFN-γ after culture with Mtb-infected macrophages as determined by ICS. The MOI was varied and the actual MOIs are shown in parentheses. Data are representative of 2 independent experiments using T cells from 5 individual mice at 4 WPI or 22 WPI (shown, A – B), analyzed in single replicates. The statistical test was a two-way ANOVA with Tukey’s post-test; actual p values are shown.

The number of intracellular bacilli could contribute to the differences in CD4 and CD8 T cell recognitions of infected macrophages. For example, a human CFP10-specific CD8 T cell clone poorly recognizes Mtb-infected DC unless they are heavily infected^24^. To determine whether the MOI affects polyclonal T cell recognition of macrophages, purified CD4 and CD8 T cells from the lungs of infected mice were cultured with macrophages infected using a range of MOI, and recognition was measured by MIM-ICS. CD4 T cells readily recognized infected-macrophages, even at a low MOI, and recognition increased at an MOI of 1.2 and plateaued at an MOI of 5.8 (Fig.5b). In contrast, there was little or no recognition by CD8 T cells at a low MOI, although recognition increased significantly when the MOI was increased to 5.8. (Fig.5b). Thus, pulmonary CD4 T cells recognize Mtb-infected macrophages with greater sensitivity than CD8 T cells.

### Quantifying T cells that recognize Mtb-infected macrophages after BCG vaccination

We hypothesize that a protective vaccine will elicit MIME^+^ T cells. We measured splenic T cell recognition of Mtb-infected macrophages after 4-5 weeks post-subcutaneous vaccination with BCG, the only approved vaccine in clinical use for TB. Under these conditions, BCG elicited T cells that recognize Mtb-infected macrophages, and more T cells recognized Mtb-infected macrophages than those that recognized the Mtb epitopes Ag85b_240-254_, 32a_93-102_, TB10.4_4-11_ or the 300 peptide megapool (Fig.6a). The dose of BCG used in our studies conferred nearly a 1 log_10_ CFU reduction when mice were challenged with Mtb nine months after vaccination (Fig.6b). This result suggests that BCG elicits a broad MIME^+^ T cell response, and this breadth in T cell recognition of infected macrophages post-vaccination might contribute to the protection conferred by BCG. Thus, the MIME could be a complementary approach to assess T cell-based vaccine candidates.

**Figure 6.**
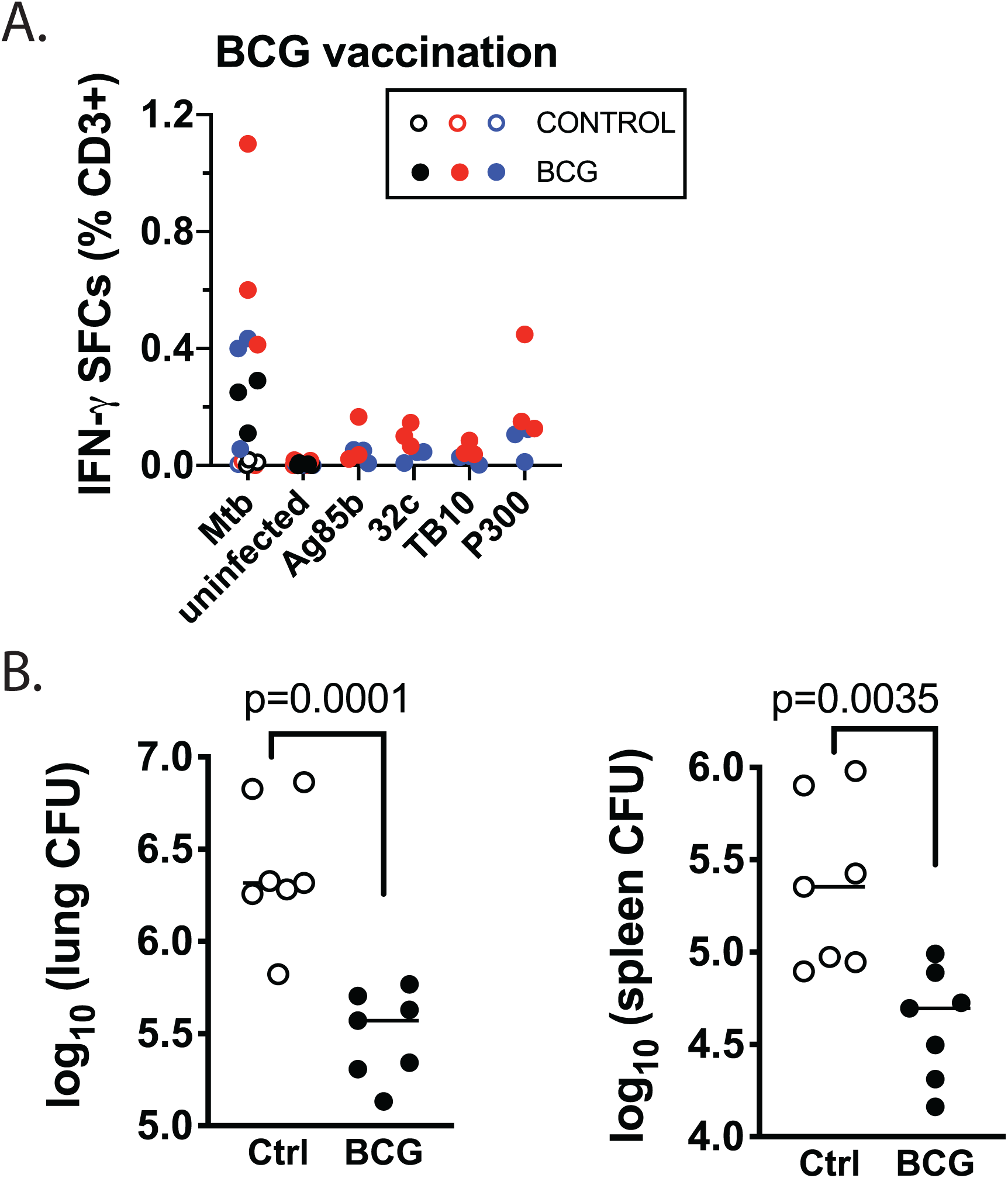
BCG elicits T cells that recognize Mtb-infected macrophages. (A) Splenic T cells were enriched by negative selection from the age-matched control (open symbols) or BCG vaccinated (closed symbols) mice, 4-5 weeks after immunization. The MIME assay was used to determine the frequency of T cells that recognized Mtb-infected macrophages. In addition, the frequency of Mtb epitope-specific T cells among these T cells was determined by coculture with uninfected macrophages and the respective peptides. Data are combined showing 3 individual mice per experiment from 2-3 experiments (color coded) (n = 9 mice for MIME response; n = 6 mice for peptide response). Each point is an average of duplicate two replicates. (B) BCG-vaccinated or control C57BL/6 mice (n=7/group) were challenged 9 months post vaccination using aerosol Mtb infection and CFU in the lungs and spleens was assessed at 4 wpi. Data from 1 representative experiment are shown.

## Discussion

Numerous microbial and host factors determine whether a protein from an intracellular bacterium elicits a T cell response. These include the bacilli’s intracellular niche, the protein’s abundance, and whether it is secreted. Much T cell-based vaccine development operates under the paradigm that Mtb-specific T cells elicited by natural infection will recognize infected APC. A corollary is that if such T cells can be elicited by vaccination, they will mediate protection against Mtb. Recent data challenge this assumption and show that not all Mtb-antigen-specific T cells recognize Mtb-infected macrophages ^4, 6, 7, 25^. This concept is consistent with data from a recent clinical trial where CD8 T cells elicited by an adenoviral-vectored vaccine expressing Mtb antigens either failed to recognize or only modestly recognized Mtb-infected DC^26^. The current paradigm needs to incorporate the possibility that some Mtb antigens, which elicit immunodominant responses, may not be presented by Mtb-infected APC^4^. A strategy to enumerate T cell recognition of Mtb-infected APC could deepen our understanding of host-pathogen interactions and assist in the development of new vaccines.

We developed the MIME to quantify T cells that recognize Mtb-infected macrophages. A majority of infected myeloid cells from the lungs contain 1 – 5 bacteria per cell ^12^. To mimic *in vivo* conditions, we used an effective median MOI of 5, which resulted in infection of 70% of the macrophages without compromising their viability. These were important considerations since bystander APC can present antigens released from Mtb-infected cells or from dying cells, as described for DC ^25, 27–30^. We hypothesized that there would be a discrepancy between the frequency of T cells that recognize peptide epitopes versus Mtb-infected macrophages. Consistent with our hypothesis, we found a minority of purified CD4 and CD8 T cells from the lungs of Mtb-infected mice recognized Mtb-infected macrophages, although the frequency was sometimes higher, particularly during chronic infection. In our studies, >90% of the CD8 T cells capable of recognizing Mtb-infected macrophages were class I MHC-restricted. The CD4 T cell response was more complicated. In the absence of class II MHC, IL-12 was sufficient to drive noncognate IFN-γ secretion by a subset of pulmonary CD4 T cells. When IL-12 was blocked, we found that majority of the IL-12 driven-IFN-γ secretion was abrogated, and >80% of the CD4 T cells were class II MHC-restricted. A similar observation was made for IL-18, which can drive antigen-experienced CD4 T cells to secrete IFN-γ following *Salmonella* infection^14, 18^. Our data show that WT Mtb-infected macrophages are recognized by CD4 T cells in the presence of a strong TCR stimulus, and this recognition is independent of IL-12.

An important finding is that, based on our IFN-γ elispot assay, the combined frequency of CD4 T cells recognizing Ag85b_240-254_ and ESAT-6_3-17_, two epitopes known to be presented by infected macrophages, accounts for only one third of the polyclonal CD4 T cells that recognize Mtb-infected macrophages. The case is even more extreme for CD8 T cells. It is unknown whether 32a_93-102_ is presented by Mtb-infected macrophages; but TB10.4_4-11_ is not ^4^. Thus, fewer than 10% of the total CD8 T cells that recognize Mtb-infected macrophages can be accounted for by known epitopes. Consistent with these observations, our MIM-ICS data strongly suggest that there are other epitopes for CD4 T cells that recognize Mtb-infected macrophages but are not represented in the multi-peptide pool 300. Conversely, there are epitopes for CD8 T cells that may not be efficiently presented by Mtb-infected macrophages but are overrepresented in the multi-peptide pool 300. The megapool was developed based on peptide recognition in humans, and thus the different MHCs in human and mice likely also contribute to differences seen, a similar peptide pool developed in mice is currently not available.

A limitation of the MIME is its focus on IFN-γ. Although IFN-γ is useful for detecting Mtb-specific T cell responses, we recognize that under some conditions or in some individuals, other cytokines (e.g., IL-2, IL-17, and TNF) may be produced by T cells in the absence of IFN-γ. Fortunately, both the ELISPOT and ICS assays can be modified to detect more than one cytokine. A second limitation is the focus on antigens that are presented during the first 48 hours of *in vitro* infection. An interesting issue was that the frequency of epitope-specific T cells differed between assays ^21, 31, 32^. Key differences in methodologies and the timing (see methods) make direct comparisons difficult. Similar discrepancies between elispot and ICS assays were previously found for the T cell response to protein antigens^32^. For the class II MHC-restricted peptide epitopes, ICS led to a greater calculated frequency than tetramers or elispot, while for the class I MHC-restricted epitopes, the frequency based on tetramer staining and ICS were similar, and greater than that determined by the elispot assay. One possibility is that the class II tetramers may inefficiently detect low affinity antigen-specific T cells, which may be more sensitively detected by dodecamers^33^. Regardless, both the Mtb-infected macrophage ELISPOT and the Mtb-infected macrophage ICS assay yield similar frequencies of IFN-γ-producing T cells that recognize Mtb-infected macrophages.

Other studies have shown that after *in vitro* expansion, human and murine CD8 T cell lines recognize and kill Mtb-infected DC or macrophages^24, 34–37^. These studies have been useful for characterizing antigen specificity and T cell function, but cannot deduce the ex vivo frequency of T cells that recognize infected macrophages. Additionally, macrophage phagosomes are more degradative compared to DC phagosomes, which may lead to differences in antigen presentation^38, 39^. Limited information is available concerning the capacity of ex vivo T cells to recognize Mtb-infected cells^24, 34–37, 40–42^. Barriers to these experiments include the low frequency of antigen-specific T cells in human blood and technical difficulties using live Mtb-infected cells, especially macrophages. Most human and murine studies use DC as the infected cell^40^. Although the Flynn lab assessed lymph node and lung T cell recognition of Mtb-infected DC, these studies found very low frequencies of T cells that recognized Mtb-infected DC^40, 43^.

Possible confounders include the use of total lung cells instead of purified T cells and the reliance on anti-MHCI or anti-MHCII antibodies to estimate the frequencies of CD4 and CD8 T cells recognizing Mtb-infected DC, respectively. The interpretation of data using class II MHC blocking antibodies is problematic because of noncognate activation of CD4 T cells, as we observed (Fig.3). While the Lewinsohn lab found that human CD4 and CD8 T cells recognize Mtb infected DC, and CD8 T cells recognize heavily infected DC, T cell clones were used for this study. Since macrophages play an important role in Mtb biology and disease progression^44–46^, we wanted to assess T cell recognition of Mtb-infected macrophages. Using a tractable system that allows the use of Mtb-infected macrophages at a median MOI of ~5, we report the ex vivo frequencies of primary T cells that recognize Mtb-infected macrophages post-infection and post-BCG vaccination.

At the present time, we have no data to directly correlate MIME^+^ T cells with control of CFU in vivo, as we do in vitro^4^. Since direct CD4 T cell recognition of infected APC is required for CFU control in vivo^8^, it is likely that MIME^+^ CD4 and CD8 T cells recognize Mtb-infected APC and promote bacillary control in vivo. Why then, despite an apparent increase in the frequency MIME^+^ T cells late during infection, is there a failure to control the lung bacterial burden? Such T cells may become dysfunctional^47^ or fail to be optimally positioned to interact with infected APC^48, 49^. Another possibility is that infected macrophages inefficiently present bacterial antigens to T cells, either because of active immune evasion or APC dysfunction. Alternatively, antigen presentation by Mtb infected cells could be limited because when intracellular bacilli are present at low numbers (i.e., <5 bacilli/cell) or are metabolically quiescent, there is little antigen available for presentation. Our data that polyclonal CD8 T cells preferentially recognize Mtb-infected macrophages at higher MOIs are consistent with published data showing human CD8 T cell clones preferentially recognize heavily infected DC^24^. Furthermore, we show that CD4 T cells are better able to recognize Mtb-infected macrophages at a higher MOI and their ability is also limited at lower MOIs. Whether this disparity could also be affected by differences in macrophages vs. DC, mouse vs. human APCs, or ex vivo vs. cultured T cells, remains to be determined. It would be interesting to characterize CD4 and CD8 T cells that can recognize low-MOI infected macrophages as this may identify antigens that are more likely to be presented during natural infection or post-challenge. Increasingly, we favor the idea that vaccine failure has little to do with the quality of the T cells that are elicited; instead, the problem is that vaccine-elicited T cells may not recognize infected APC that harbor few bacilli.

The low frequency of MIME^+^ T cells is consistent with Mtb using multiple strategies to evade detection by the immune system ^50^. We and others have proposed that Mtb could be using certain immunodominant antigens as decoys ^4, 51^. An understanding of the host-pathogen interactions will be crucial in identifying protective antigens for use in the next generation of vaccines. We show that T cells elicited by BCG vaccination are capable of recognizing Mtb-infected macrophages. The frequency of BCG-elicited T cells could be used as a benchmark to compare other whole cell or subunit vaccines. A correlation between T cells recognizing Mtb-infected cells and protection could also provide insights into the immunological mechanisms of novel vaccines.

We envision that the MIME assay could be used to study immune evasion by Mtb and study T cell recognition of Mtb-infected macrophages. For example, we previously used a modified version of the MIME assay to show that iNKT cells recognized Mtb-infected macrophages, and the MIME assay could be applied to other T cell subsets (e.g., unconventional T cells) ^52^. Finally, as we believe that recognition of Mtb-infected macrophages by vaccine-elicited T cells will be a prerequisite to protection, we expect that the MIME assay could be used to assess T cell-based vaccine candidates as a complementary approach to immunogenicity studies or other approaches such as the mycobacterial growth inhibition assay.

## Materials and Methods

### Ethics Statement

All animal studies were conducted in accordance with the protocol approved by the Institutional Animal Care and Use Committee at the University of Massachusetts Medical School (Animal Welfare Assurance number A3306-01). All studies adhere to the relevant guidelines and recommendations from the Guide for the Care and Use of Laboratory Animals of the National Institutes of Health and the Office of Laboratory Animal Welfare.

### Mice

C57BL/6J mice were purchased from The Jackson Laboratory (Bar Harbor, ME). B6.129P2-*H2-K1^tm1Bpe^H2-D1^tm1Bpe^/*DcrJ, i.e., K^b−/−^D^b−/−^ (MHC-I^−/−^) mice, originally purchased from the Jackson Laboratory, were a generous gift from Dr. Kenneth Rock (University of Massachusetts Medical School, MA). B6.129*-H2-Ab1^tm1Gru^*N12 (MHC-II^−/−^) and control C57BL/6NTac mice were purchased from Taconic Biosciences (Rensselaer, NY). C7 TCR transgenic (C7) mice were a generous gift from Dr. Eric Pamer (Memorial Sloan Kettering Cancer Center, NY)^53^.

### *In vivo* aerosol infection

Low-dose Mtb strain Erdman aerosol infections were performed as described previously^22^. Briefly, a frozen bacterial aliquot was thawed, sonicated for 1 minute using a cup-horn sonicator and diluted in 0.9% NaCl–0.02% Tween-80 in a total volume of 5 ml used for nebulization. Infections were performed using a Glas-Col (Terre Haute, IN) full body inhalation exposure system. Mice received an inoculation dose of 25-100 CFU/mouse as determined by plating undiluted lung homogenates on 7H11 plates from a subset of infected mice within 24 hours post-infection.

### Bacteria

For *in vitro* infections, Mtb strain H37Rv or H37Rv expressing YFP (H37Rv-YFP) was grown as previously described ^52, 54^. Bacteria was grown to a log phase under OD_600_ = 1.0, opsonized with TB coat (RPMI 1640, 1% heat-inactivated FBS, 2% human serum, 0.05% Tween-80), washed again and filtered through a 5 μm filter. Bacteria was counted using a Petroff-Hausser chamber before infection. H37Rv-YFP was a generous gift from Dr. Christopher Sassetti (University of Massachusetts Medical School, MA).

### *In vitro* Mtb infection

Macrophages (10^6^/well) were cultured overnight at 37°C, 5% CO_2_ in 12-well Nunc UpCell plates. Unless otherwise mentioned, H37Rv was used for infection at a multiplicity of infection (MOI) of 4. Bacteria were added to macrophages for 18-24 h at 37°C, 5% CO_2_. Then, the plates were left at room temperature for 30 minutes to allow the macrophages to become nonadherent. Macrophages were harvested by gently pipetting the cells, followed by washing each well twice. The harvested cells were washed twice by centrifugation at 1500 rpm for 5 minutes. Live, Mtb-infected or uninfected macrophages were counted by Trypan blue exclusion of dead cells and used in MIME. To assess actual MOI, a subset of the macrophages was lysed using a final concentration of 1% Triton-X-100, serially diluted using 0.9% NaCl–0.02% Tween-80 and immediately plated on 7H11 plates.

### Mtb-infected macrophage ELISPO

Mtb-infected or uninfected macrophages resuspended in complete media without antibiotics were aliquoted at 10^5^ cells/well in an elispot plate that had been coated with the IFN-γ capture antibody and blocked with complete media as per manufacturer’s instructions. Mtb-infected macrophages were allowed to adhere for at least 1 h at 37°C, 5% CO_2_, and then T cells were added to the macrophages at variable T cell:APC ratio. Positive controls included and anti-CD3 and anti-CD28 condition, (each at 2 μg/ml). Where indicated, single peptides (10 μM) or the peptide megapool 300 (2 μg/ml) were added to uninfected macrophages prior to the addition of T cells. T cells were coincubated with macrophages for 18-24 h, followed by cell lysis using deionized water, incubation with detection antibody and development as per manufacturer’s instructions. The elispot insert was fixed with 1% paraformaldehyde in PBS for 1 h and then washed 3x with water. The ELISPOT insert was allowed to dry overnight, and single-color red spots were enumerated using the CTL ImmunoSpot S5 Analyzer, software version 5.4.0.10 from Cellular Technology Limited (Cleveland, Ohio). Spot forming cells were counted and corrected for background by subtracting the spots for the conditions with infected or uninfected macrophages without any added T cells ^31^.

### Flow cytometric staining and intracellular cytokine staining (ICS)

Mtb-infected macrophages were stained with Zombie Aqua (live/dead staining) as per manufacturer’s instructions, followed by Fc receptor block with monoclonal antibody 2.4G2 and surface staining using anti-F4/80 and anti-CD45. For MHC class II tetramer staining, enriched T cells were resuspended in complete media and incubated at 37°C, 5% CO_2_ for 1 hour with MHC class II tetramers, following which surface staining was performed with antibodies at 4°C. For MHC class I tetramer staining, cells were surface stained together with tetramers and antibodies at 4°C. For experiments involving ICS, T cells enriched from Mtb-infected lung samples were cocultured with Mtb-infected macrophages, uninfected macrophages or uninfected macrophages pulsed with single peptides (10 μM) in complete media for 1 h at 37°C, 5% CO_2_. After this initial incubation, GolgiStop was added for 4 h at 37°C, 5% CO_2_ and cells were surface stained with antibodies, followed by permeabilization and staining for IFN-γ as per manufacturer’s instructions. All cells were fixed with 1% PFA before flow cytometric analysis. For the megapool 300 experiment, lung cells were cultured with Mtb-infected macrophages, uninfected macrophages, or uninfected macrophages plus megapool 300 (1 μg/ml) followed by ICS as described above. Samples were acquired using MACSQuant (Miltenyi Biotec, Germany) and analyzed using FlowJo version 9.0 (Ashland, OR).

## Acknowledgements

We thank members of the Behar lab and Kim West (University of Massachusetts Medical School) for technical assistance and discussion. We thank Dr. Christopher Sassetti, Dr. Kadamaba Papavinasasundaram, Megan Proulx and Dr. Kenneth Rock (University of Massachusetts) for reagents, assistance and discussion. We would like to thank the National Institutes of Health Tetramer Core Facility for providing reagents. Supported by R21 AI136922 and R01 AI106725 (SMB).

## Supplementary Information

### Supplementary Methods

#### Materials

The mouse interferon-γ (IFN-γ) ELISPOT kit and AEC substrate were purchased from BD Biosciences (San Jose, CA). The following peptides used for vaccination or *in vitro* experiments were purchased from New England Peptides (Gardner, MA): TB10.4_4-11_ (IMYNYPAM), Mtb32_93-102_ (GAPINSATAM), Ag85b_240-254_ (FQDAYNAAGGHNAVF) and ESAT-6_3-17_ (EQQWNFAGIEAAASA). Mtb peptide megapool 300 was previously described^23^. The MojoSort negative selection kits for CD4 and CD8 enrichment were purchased from Biolegend (San Diego, CA). Zombie Aqua viability dye, LEAF purified anti-mouse CD3ε (clone 145-2C11), LEAF purified anti-mouse CD28 (clone 37.51), LEAF purified anti-mouse IL-12 (clone C17.8), LEAF purified rat IgG2a, k (clone RTK 2758) LEAF purified rat IgG1, k (clone RTK2071), and all fluorophore conjugated antibodies used for flow cytometry were purchased from Biolegend, i.e., anti-CD4 (clone-GK1.5), anti-CD8α (clone 53-6.7), anti-CD3ε (clone 145-2C11), anti-CD19 (clone 6D5), anti-CD45 (clone 30-F11), anti-F4/80 (clone BM8), anti-CD69 (clone H1.2F3) and anti-IFN-γ (clone XMG-1.2). RPMI 1640, HEPES, sodium pyruvate and L-glutamine were purchased from Invitrogen Life Technologies, ThermoFisher (Waltham, MA). Heat inactivated fetal bovine serum was purchased from GE Healthcare Life Sciences (Pittsburgh, PA). Nunc UpCell plates were purchased from ThermoFisher Scientific (Waltham, MA). Collagenase type IV was purchased from Sigma-Aldrich (St. Louis, MO). CD4 (L3T4), CD8a (Ly-2), CD90.2 and CD11b microbeads used for positive selection were purchased from Miltenyi Biotec (Germany). Peptide-loaded tetramers for ESAT-6_3-_17 (I-A^b^), Ag85b_240-254_ (I-A^b^), Mtb32_93-102_ (H2-D^b^), TB10.4_4-11_ (H2-K^b^) were obtained from the National Institutes of Health Tetramer Core Facility (Emory University Vaccine Center, Atlanta, GA). Corning human AB serum, Triton-X-100 and Tween-80 were purchased from Fisher Scientific. GolgiStop and BD Cytofix/Cytoperm were purchased from BD Pharmingen (San Jose, CA). 7H11 plates were purchased from Hardy Diagnostics (Santa Maria, CA).

#### Macrophages

For isolation of murine thioglycollate-induced peritoneal macrophages (TG-PM), naïve mice were intraperitoneally injected with 2 ml of 3% thioglycollate solution. Macrophages were harvested by peritoneal lavage 4-5 days. Macrophages were CD11b-enriched as per manufacturer’s instructions. Enriched macrophages were resuspended in complete media without antibiotics (RPMI 1640 supplemented with 10% fetal bovine serum, 10 mM HEPES, 1 mM sodium pyruvate and 2 mM L-glutamine) and used for Mtb-infected macrophage ELISPOT or other assays as described. Purity of the enriched macrophages was confirmed by surface-staining of cells and cells were greater than 90% CD45^+^F4/80^+^.

#### T-cell isolation from infected or vaccinated mice

For experiments involving pulmonary T cells, lungs were dissected from infected mice after perfusion with RPMI 1640. Single cell lung suspensions were prepared by coarse dissociation using the GentleMACS tissue dissociator (Miltenyi Biotec, Germany), followed by digestion for 30 min at 37°C in a shaker at 85 rpm with 300 U/ml of Collagenase type IV in complete media. Samples were processed again using the GentleMACS tissue dissociator, strained through a 70 μm filter, washed 1x with PBS and then strained through a 40 μm filter. CD4 or CD8 T cells were labeled with CD4 L3T4 microbeads or CD8a Ly-2 microbeads, respectively, followed by positive selection though magnetic column using AutoMACS (Miltenyi Biotec, Germany). Where indicated, CD90.2 beads were used for CD4 and CD8 T cell positive selection. T cells were resuspended in complete media without antibiotics before use. Purity of the enriched T cells was confirmed by surface staining for CD4, CD8 and CD3 and was greater than 90%.

For experiments involving splenic T cells, spleens from infected or vaccinated mice were dissociated using a syringe and filtered through a 70 μm filter. Splenocytes were washed 1x with PBS and strained through a 40 μm filter. Polyclonal CD4 or bulk T cells (CD4 and CD8 T cells) were enriched using Mojosort negative selection kits for CD4 and CD3, respectively. T cells were resuspended in complete media without antibiotics before use. Purity of the enriched T cells was confirmed by surface staining and was greater than 90%.

#### T-cell lines

C7 CD4^+^ T cells (specific for ESAT-6) and P25 CD4^+^ T cells (specific for Ag85b) have been described previously^4^. Briefly, C7 or P25 cell lines were stimulated *in vitro* with irradiated splenocytes pulsed with 10 μM peptide ESAT-6_3-17_ (EQQWNFAGIEAAASA) or Ag85b_240-254_ (FQDAYNAAGGHNAVF) in complete media containing IL-2 at 20 u/ml, in the absence of antibiotics. After the initial stimulation, the T-cell cultures were split every two days for 3-4 divisions and rested for at least three weeks post-initial stimulation before use. After the initial stimulation, the cells were cultured in complete media containing IL-2 and IL-7 and final concentrations of 20 U/ml and 10 ng/ml, respectively. T cells were used between three – six weeks post-initial stimulation. Purity of C7 T cell line was assessed by Vb10.

#### BCG vaccination

Mice were vaccinated subcutaneously with a single dose of BCG strain SSI diluted in 0.04% Tween-80 in PBS. BCG strain SSI was generously provided by Dr. Christopher Sassetti (University of Massachusetts Medical School). The BCG stocks used for vaccination were previously frozen and thawed immediately before use, washed 2x with 0.04% Tween-80 in PBS and used at an average dose of 500,000 CFU/mouse in a final volume of 200 μl. To confirm the dose, bacteria used for vaccination were plated on 7H10 plates. Splenic T cells were enriched at 4-5 weeks and used for the Mtb-infected macrophage ELISPOT as described above.

### Supplementary Figures

**Supplementary Figure 1.**
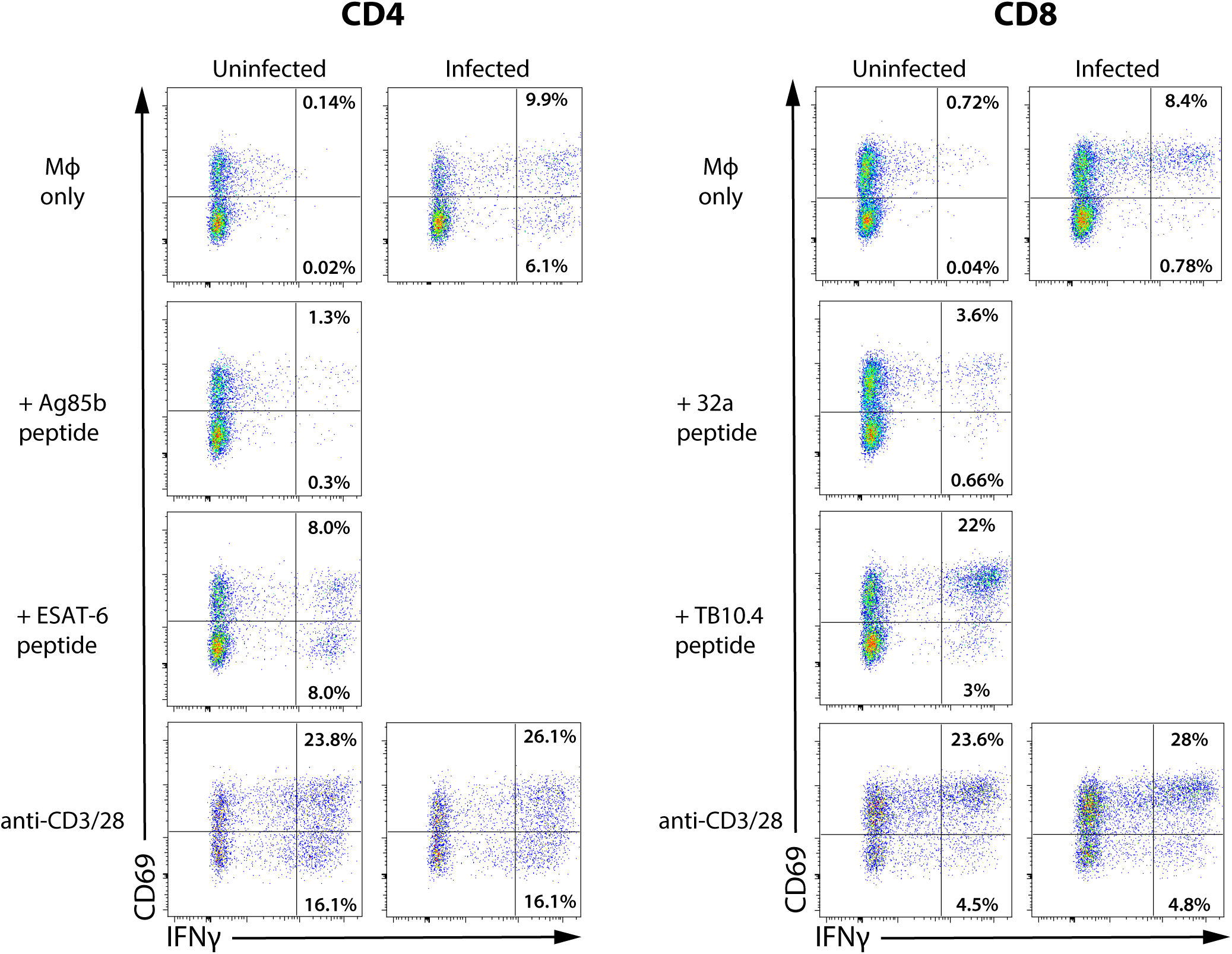
The majority of T cells that recognize Mtb-infected macrophages express CD69^+^. Pulmonary CD8 and CD4 T cells were cocultured with uninfected macrophages with the respective peptides or infected macrophages as indicated and assessed for IFN-γ production by ICS at 5 hours (D, E). Data are representative of 2 independent experiments using pooled T cells from 5 mice at 5 WPI (shown) or 7.5 months post infection, analyzed in triplicates.

